# Selective constraints on coding sequences of nervous system genes are a major determinant of duplicate gene retention in vertebrates

**DOI:** 10.1101/072959

**Authors:** Julien Roux, Jialin Liu, Marc Robinson-Rechavi

**Affiliations:** Université de Lausanne, Département d’Ecologie et d’Evolution, Quartier Sorge, 1015 Lausanne, Switzerland; Swiss Institute of Bioinformatics, Lausanne, Switzerland.

**Keywords:** Whole-genome duplication, small-scale duplication, neuron, anatomy, gene expression, protein misfolding, protein interaction, translational accuracy

## Abstract

The evolutionary history of vertebrates is marked by three ancient whole-genome duplications: two successive rounds in the ancestor of vertebrates, and a third one specific to teleost fishes. Biased loss of most duplicates enriched the genome for specific genes, such as slow evolving genes, but this selective retention process is not well understood. To understand what drives the long-term preservation of duplicate genes, we characterized duplicated genes in terms of their expression patterns. We used a new method of expression enrichment analysis, TopAnat, applied to *in situ* hybridization data from thousands of genes from zebrafish and mouse. We showed that the presence of expression in the nervous system is a good predictor of a higher rate of retention of duplicate genes after whole-genome duplication. Further analyses suggest that purifying selection against the toxic effects of misfolded or misinteracting proteins, which is particularly strong in non-renewing neural tissues, likely constrains the evolution of coding sequences of nervous system genes, leading indirectly to the preservation of duplicate genes after whole-genome duplication. Whole-genome duplications thus greatly contributed to the expansion of the toolkit of genes available for the evolution of profound novelties of the nervous system at the base of the vertebrate radiation.

## Introduction

The process of gene duplication plays a major role in the evolution of genomes, as it provides raw material for innovation (Lynch and Conery 2000; Conant and Wolfe 2008; Van de Peer et al. 2009). But only a minority of the gene duplication events reach fixation in a species, and survive in the long term with two functional gene copies (Innan and Kondrashov 2010). It is not yet clear what factors drive this process of selective retention, but it is clear that it is non random (Davis and Petrov 2004).

Focusing on whole-genome duplication events allows quantification of the long-term retention bias alone, the whole gene set having been fixed in duplicate (Singh et al. 2012). In vertebrates, it has been estimated that only 10 to 20% of the duplicates (or “ohnologs”) that originated from the ancient whole-genome duplications at the origin of the lineage (“2R” hypothesis) (Ohno 1970; Holland et al. 1994; Hughes 1999; Putnam et al. 2008) or in teleost fishes (“3R” hypothesis) (Jaillon et al. 2004; Meyer and Van de Peer 2005) were eventually retained in the long term (Brunet et al. 2006; Nakatani et al. 2007; Putnam et al. 2008; Smith and Keinath 2015). Retained genes do not constitute a random subset of genes. For instance, their protein sequences tend to be under strong selective constraint (Davis and Petrov 2004; Brunet et al. 2006; Howe et al. 2013). They tend to be involved in functions such as signaling, cognition and behavior, or regulation of transcription (Brunet et al. 2006; Putnam et al. 2008; Kassahn et al. 2009; Huminiecki and Heldin 2010; Schartl et al. 2013), and to be expressed late in development (Roux and Robinson-Rechavi 2008). The causal mechanisms linking such properties to increased retention after whole-genome duplication have not been fully clarified so far.

An interesting study found that in yeast and *Paramecium*, the expression level of genes was a major determinant of their duplication retention rate after whole-genome duplication (Gout et al. 2010), highly expressed genes being more retained, an effect that could not be explained indirectly by other factors. This observation is noteworthy since gene expression level is also known to be a major determinant of the rate of protein evolution across a wide range of species (Drummond et al. 2005; Gu and Su 2007; Drummond and Wilke 2008), highly expressed genes having lower rates of protein evolution.

The generalization of this result to vertebrates is complicated by their complex anatomy. One way to address this complexity is to investigate whether patterns of expression over anatomy could be linked to ohnolog retention rates. But surprisingly this question has rarely been addressed. Based on EST data, a study found little association between expression breadth – the number of tissues in which a gene is expressed – and retention rate after the 2R whole-genome duplication (Satake et al. 2012). However, the authors observed lower retention of the fast-evolving genes expressed in endodermal tissues, such as the digestive tract, compared to slow-evolving genes expressed in ectodermal tissues, such as the nervous system. The expression patterns were opposite for small-scale duplication events, in agreement with other results showing that these two types of duplications tend to affect opposite sets of genes (Davis and Petrov 2005; Makino et al. 2009).

These observations suggest that anatomical expression patterns of genes might help to understand the process of ohnolog retention in vertebrates. Unfortunately the techniques used to study gene expression patterns on a genomic scale, previously ESTs and microarrays, and more recently RNA-seq, usually lack anatomical precision. In this paper we took advantage of bioinformatics integration of another source of expression data, *in-situ* hybridizations. Expression patterns obtained with this technique are very precise, sometimes down to the cellular resolution (Lein et al. 2007; Diez-Roux et al. 2011; Jacobs et al. 2011). They are also very inclusive, since it is possible to visualize the expression of a particular gene in the entirety of anatomical structures present in a histological section or even an entire organism (“whole-mount” *in-situ* hybridizations), without selecting *a priori* a tissue to dissect. Compared to other techniques, there is also less averaging or dilution of the expression signal for genes whose expression is heterogeneous among the cells or substructures of a tissue (Altschuler and Wu 2010; Pantalacci and Semon 2015).

A drawback of *in-situ* hybridizations, however, is that they usually give information on only one, or sometimes a handful of genes. Fortunately, there have been several efforts to generate with this technique high-throughput atlases of gene expression patterns in model organisms, notably zebrafish and mouse (e.g., Neidhardt et al. 2000; Thisse et al. 2004; Lein et al. 2007; Diez-Roux et al. 2011). Thus, there are thousands of *in-situ* hybridizations publicly available, allowing us to perform analyses at the genomic scale. Even more valuable, the expression patterns revealed by these hybridizations have been manually annotated to terms from anatomical ontologies, notably the cross-species ontology Uberon describing anatomical structures and their relationships in animals (Hayamizu et al. 2005; Sprague et al. 2006; Bastian et al. 2008; Mungall et al. 2012; Hayamizu et al. 2013; Haendel et al. 2014).

To detect the biases in anatomical expression patterns of ohnologs, we developed a novel bioinformatics approach. Similarly to the widely used functional enrichment tests performed on categories of the Gene Ontology (Ashburner et al. 2000; Yon Rhee et al. 2008), we used a Fisher’s exact test to detect an enrichment in the proportion of ohnologs expressed in each anatomical structure of the organism. This methodology allowed us to monitor expression biases with great precision, and to benefit from the information encoded in the ontology (e.g., parent-child relationships).

We observed that genes expressed in the nervous system had an increased chance of being retained after whole-genome duplication, whereas they had a decreased chance of being duplicated via small-scale duplication. This novel and robust observation helped us clarify the gene properties that causally influence the retention of duplicate genes. The rate of non-synonymous substitutions of nervous-system genes, their level of optimization of synonymous codon usage at sites that are important for protein structure stability, and their maximum level of expression across neural tissues are significantly associated to retention rate, suggesting a major role of purifying selection on coding sequence on ohnolog retention in vertebrates. This selective force is particularly strong on nervous system genes, primarily preventing them from producing toxic protein products. It could have the unexpected consequence of lowering their probability of loss of function, leading to their evolutionary long-term retention.

This model is consistent with a model proposed to explain the counterintuitive expansion of human disease-causing genes after the 2R whole-genome duplication events (Singh et al. 2012; Malaguti et al. 2014), and is not exclusive of previously proposed models, e.g., sub- or neo-functionalization (Ohno 1970; Force et al. 1999; He and Zhang 2005), the dosage-balance hypothesis (Freeling and Thomas 2006; Makino and McLysaght 2010; Birchler and Veitia 2012; Singh et al. 2012), or selection for absolute dosage (Osborn et al. 2003; Innan and Kondrashov 2010). Rather it expands these models to illustrate the key role of anatomy in shaping the duplicated gene content of vertebrate genomes.

## Results

### 3R ohnologs are biased for nervous system expression

Zebrafish 3R ohnologs were identified using a phylogenomics approach, and were used as input gene list in an expression enrichment test (Materials and Methods, Figure S1). The list of anatomical structures showing enrichment for expression of these genes is shown in Table 1. At a false discovery rate (FDR) threshold of 10%, 25 structures were significantly enriched. The only significant depletion was for “unspecified”, a term indicating that the gene expression was assayed, but no anatomical structure was specified by the author.

**Table 1:** Zebrafish anatomical structures showing a significant enrichment in expression of zebrafish 3R ohnologs (FDR < 10%). Anatomical structures are sorted by their enrichment fold compared to null expectation. The sub-structures of the “broad” nervous system are highlighted in bold. The “weight” algorithm of the topGO package was used to decorrelate the structure of the ontology. The full list of anatomical structures, sorted by *p*-value, is shown in Table S1.

| Organ ID | Organ name | Number of genes expressed | Number of 3R ohnologs expressed | Number of 3R ohnologs expected | Enrichment fold | *p*-value | FDR |
| --- | --- | --- | --- | --- | --- | --- | --- |
| UBERON:2007001 | **dorso-rostral cluster** | 18 | 9 | 1.88 | 4.79 | 2.91E-05 | 3.02E-03 |
| UBERON:2007002 | **ventro-rostral cluster** | 21 | 10 | 2.19 | 4.57 | 1.78E-05 | 2.00E-03 |
| UBERON:2007003 | **ventro-caudal cluster** | 19 | 9 | 1.99 | 4.52 | 5.01E-05 | 4.23E-03 |
| UBERON:0000204 | **ventral part of telencephalon** | 102 | 25 | 10.66 | 2.35 | 3.48E-05 | 3.36E-03 |
| UBERON:0002946 | **regional part of cerebellum** | 79 | 19 | 8.26 | 2.30 | 3.86E-04 | 2.74E-02 |
| UBERON:0002757 | **regional part of epithalamus** | 546 | 121 | 57.06 | 2.12 | 1.30E-16 | 8.78E-14 |
| UBERON:0008904 | **neuromast** | 169 | 37 | 17.66 | 2.10 | 8.91E-06 | 1.09E-03 |
| UBERON:0000203 | **pallium** | 105 | 23 | 10.97 | 2.10 | 4.27E-04 | 2.88E-02 |
| UBERON:0003895 | hypaxial myotome region | 101 | 22 | 10.55 | 2.09 | 6.13E-04 | 3.76E-02 |
| UBERON:0010134 | **secretory circumventricular organ** | 478 | 104 | 49.95 | 2.08 | 8.10E-14 | 3.64E-11 |
| UBERON:0003902 | **retinal neural layer** | 633 | 136 | 66.15 | 2.06 | 2.09E-17 | 2.82E-14 |
| UBERON:0003296 | **gland of diencephalon** | 559 | 115 | 58.42 | 1.97 | 1.70E-13 | 5.73E-11 |
| UBERON:0002540 | **lateral line system** | 519 | 101 | 54.24 | 1.86 | 6.78E-06 | 9.15E-04 |
| UBERON:0001898 | **hypothalamus** | 329 | 59 | 34.38 | 1.72 | 6.10E-04 | 3.76E-02 |
| UBERON:0000045 | **ganglion** | 596 | 105 | 62.28 | 1.69 | 2.26E-08 | 5.07E-06 |
| UBERON:0002199 | integument | 565 | 97 | 59.04 | 1.64 | 3.51E-07 | 5.92E-05 |
| UBERON:0001894 | **diencephalon** | 1459 | 239 | 152.47 | 1.57 | 1.82E-03 | 9.82E-02 |
| UBERON:0005725 | **olfactory system** | 760 | 124 | 79.42 | 1.56 | 6.94E-08 | 1.34E-05 |
| UBERON:0002298 | **brainstem** | 624 | 101 | 65.21 | 1.55 | 3.79E-05 | 3.41E-03 |
| UBERON:0003051 | ear vesicle | 843 | 133 | 88.09 | 1.51 | 9.98E-07 | 1.50E-04 |
| UBERON:0000489 | cavitated compound organ | 1701 | 255 | 177.76 | 1.43 | 1.34E-08 | 3.63E-06 |
| UBERON:0002028 | **hindbrain** | 1600 | 225 | 167.2 | 1.35 | 7.71E-05 | 6.11E-03 |
| UBERON:0000479 | tissue | 4751 | 645 | 496.49 | 1.30 | 1.29E-03 | 7.27E-02 |
| UBERON:0000955 | **brain** | 3255 | 435 | 340.15 | 1.28 | 1.29E-04 | 9.70E-03 |
| UBERON:0000483 | epithelium | 3734 | 489 | 390.21 | 1.25 | 8.56E-04 | 5.02E-02 |

A high fraction of the enriched anatomical structures were neural (e.g., sub-parts of the telencephalon, cerebellum, epithalamus, neuromast, retinal neural layer). To test whether this did not simply reflect the structure of the Uberon anatomical ontology, which could use more terms to describe the nervous system than other anatomical systems, we made the inventory of all nervous system structures described in the ontology. We built two lists, a strictly defined list, and a broader one also including sensory systems as well as embryonic precursors of nervous structures (“strict” and “broad” lists; see Materials and Methods). Using these reference lists we observed that among the 25 structures shown to be significantly enriched for the expression of 3R ohnologs, 19 were part of the nervous system (broad list; of which 15 were part of the strict list). These proportions were significantly higher than the proportion of nervous system structures among all tested structures (Fisher’s exact tests; broad list: *p*=0.0001, with odds-ratio=5.41; strict list: *p*=3.2e-5, odds-ratio=5.67). Even among structures that were not part of the nervous system, some still shared the same ectoderm developmental origin as nervous system (e.g., “ear vesicle” or “integument”).

We applied the same procedure to other anatomical systems (see Materials and Methods), but they were always under-represented among the structures enriched for the expression of 3R ohnologs. We also verified that the over-representation of nervous system structures was not dependent on the FDR threshold used in the enrichment analysis. In the rest of the article we describe the ontology enrichment results obtained at a FDR threshold of 10%, and we use the broad list of nervous system structures as reference.

We next turned to singleton genes, whose duplicate copy was lost after the 3R whole-genome duplication, and investigated if these were preferentially expressed in any anatomical structure. We found only two structures enriched for this group of genes: “unspecified” and “alar plate midbrain”. However, 35 structures were significantly depleted in expression of singletons (Table S2), of which 22 were part of the nervous system (Fisher’s exact test; *p*=0.0024, odds=2.89)

In summary, we observed that genes retained in duplicate after the fish-specific (“3R”) whole-genome duplication were strongly biased for nervous system expression (in very diverse structures, including developmental precursors and sensory organs), whereas genes that were not retained in duplicate had the opposite tendency to not show expression in these structures. We reproduced these analyses using an independent dataset of 3,212 and 10,415 zebrafish 3R ohnologs and singletons identified using phylogenetic and synteny analyses (Braasch et al. 2016), and obtained consistent results (Tables S3 and S4).

### Pre- or post-duplication bias?

These results could be explained by a duplicate retention bias, i.e., genes expressed in the nervous system before 3R were more likely retained as ohnologs. Or they could be explained by a bias in post-duplication evolution, i.e., ohnologs were more likely to acquire expression in the nervous system. To disentangle these two scenarios, it is possible to focus on an outgroup species that did not experience the whole-genome duplication, and compare the properties of orthologs of ohnologs to orthologs of singletons in this species, as a proxy for the pre-duplication properties of zebrafish genes (Davis and Petrov 2004; Brunet et al. 2006; Roux and Robinson-Rechavi 2008). The mouse represents a convenient such outgroup, since a large number of *in-situ* hybridization data are also available for this species, allowing to test the enrichment of expression in anatomical structures using the same methodology (see Materials and Methods).

We found that mouse orthologs of zebrafish 3R ohnologs were enriched for expression in 57 anatomical structures, among which 46 were nervous system structures (Table S5; *p*=1.6e-19, odds=13.9). In parallel, mouse orthologs of 3R singletons were significantly depleted for expression in two nervous structures, “olfactory cortex mantle layer” and “CA2 field of hippocampus”, and just above significance threshold, nervous system structures were also almost exclusively present at the top of the list (Table S6).

These results, consistent with the observations in zebrafish, suggest that the nervous system enrichment can be explained by an ohnolog retention bias, and that expression patterns before the 3R whole-genome duplication, or in an outgroup, can predict this retention bias.

### The nervous system bias is weakly detected for 2R ohnologs

We repeated the enrichment analysis with mouse 2R ohnologs identified by phylogenomics, but these genes did not show any significant enrichment (Table S7). However, there was a significant enrichment for nervous system structures when we used an independent list of 5,376 mouse 2R ohnologs (Singh et al. 2015), identified using synteny comparison across multiple genomes (Table S8; 91 nervous structures out of 297 enriched structures; *p*=0.0081, odds=1.44).

Mouse 2R singletons were depleted in 107 structures, 31 of which belonged to the nervous system (Table S9). This slight over-representation of nervous system structures was however not significant (*p*=0.25, odds=1.29).

In summary, the results from the 2R whole-genome duplications were consistent with those from the 3R whole-genome duplication. Several technical factors could account for the fact that the 2R trends were weaker than the 3R trends, notably the older age of these events, but also post-duplication evolution patterns confounding this analysis, that was not performed in an outgroup species.

### Small-scale duplications

We also investigated whether an anatomical expression bias existed for duplicate genes that arose from other sources than whole-genome duplications, i.e., small-scale duplications. Because there was no whole-genome duplication in the phylogenetic branches leading to the zebrafish and mouse lineages, after 3R and 2R respectively, we isolated duplicates dated to these branches as small-scale duplicates. We removed those that were specific to these species since they could still be polymorphic or represent errors in the genome assemblies (Materials and Methods). Unfortunately, the small number of genes identified (385 and 646 duplicate genes for zebrafish and mouse respectively) led to low power of the enrichment test. In both species, we did not detect any significantly enriched or depleted anatomical structure. In mouse, the depletion results were close to significance, and we noticed numerous nervous system structures present at the top of the list (Table S10). Using an external curated list of small-scale duplicate pairs specific to rodents (Farre and Alba 2010), there were four structures with a significant expression enrichment (“placenta”, “stomach glandular region mucosa”, “ectoplacental cone” and “cardia of stomach”) and 8 structures with a significant depletion, 7 of which were part of the nervous system (*p*=0.0003, odds=22.1; Table S11). Overall, there was weak evidence for a nervous system expression bias of small-scale duplicates.

### Validation with microarray data

To check whether the expression biases could be observed with other types of expression data, we retrieved a microarray dataset in mouse that included samples from multiple nervous and non-nervous tissues (see Materials and Methods). We called the genes expressed or not in each sample, and ranked the tissues based on the proportion of mouse orthologs of zebrafish 3R ohnologs expressed (Figure 1A). We observed that the samples expressing the highest proportion of orthologs of ohnologs belonged to the nervous system. This result was confirmed with another microarray dataset (Figure S2A), and with human RNA-seq data from the GTEx consortium (Figure S2B).

**Figure 1:**
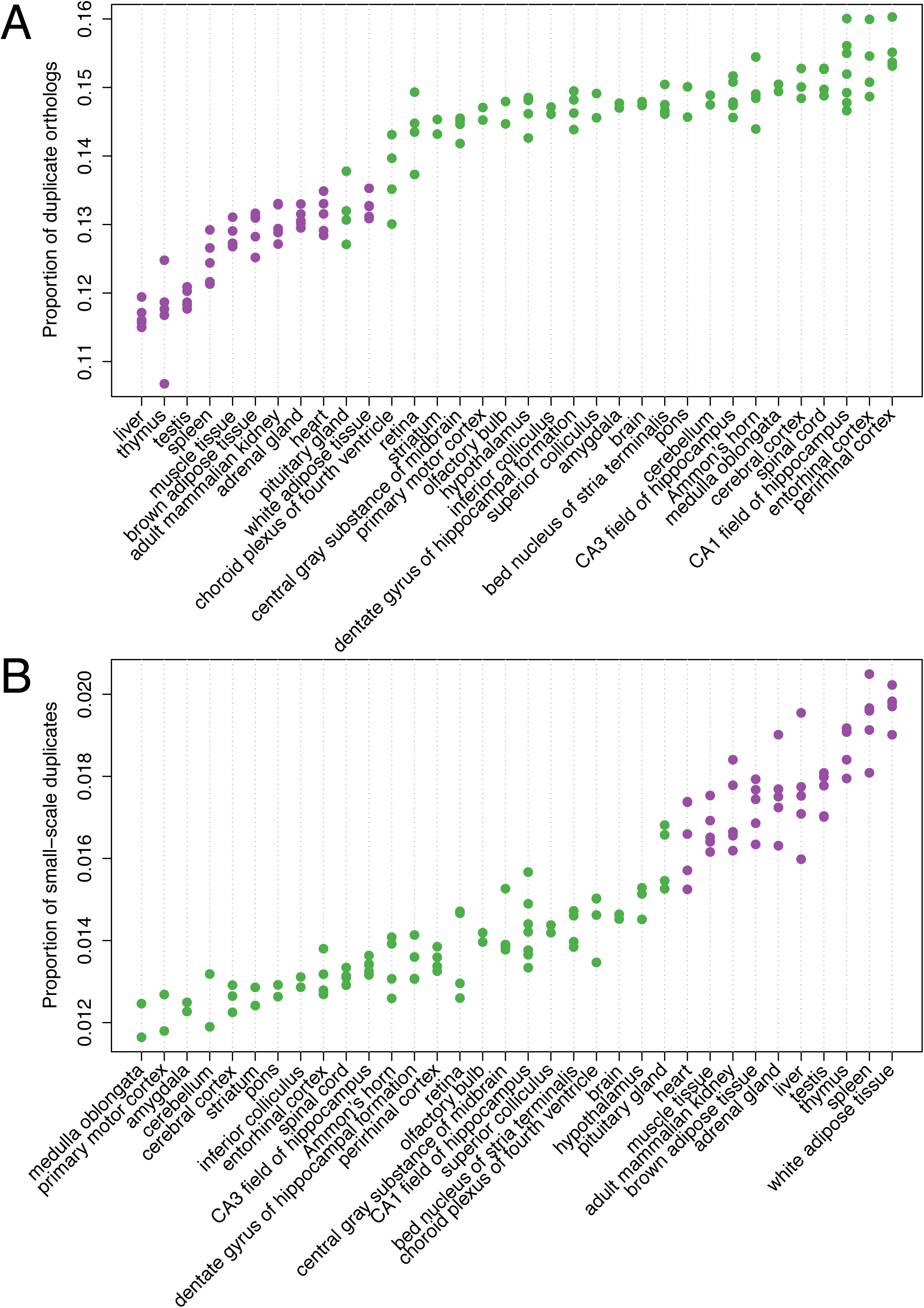
Proportion of mouse orthologs of zebrafish 3R ohnologs among genes expressed in the different tissues sampled in the GSE3594 microarray experiment. The reference gene set was composed only of mouse orthologs of zebrafish 3R ohnologs and singletons. Tissues are ranked based on the average proportion of orthologs of ohnologs expressed. Each dot represents a sample (biological replicate). Green color represents nervous-system tissues and purple represents non-nervous-system tissues.

Across neural tissues, there was little variation in the proportion of orthologs of ohnologs expressed, confirming that this bias was general to the whole nervous system. Only a few neural tissues stood out with lower proportions, notably the pituitary gland (Figure 1A, S2A and S2B). An independent microarray dataset, including samples from 46 neural tissues confirmed that the pituitary gland, but also the pineal body, displayed a lower proportion of orthologs of ohnologs expressed (Figure S2C). Interestingly, these tissues also stood out from clustering analyses based on expression levels across numerous nervous tissues (Zapala et al. 2005; Kasukawa et al. 2011), possibly because of their secretory activities (Gu and Su 2007) or different cell type composition.

Finally we ranked tissues based on the proportion of small-scale duplicates expressed, and observed an opposite picture: tissues expressing the lowest proportion of small-scale duplicates belonged to the nervous system (Figure 1B), supporting the weak trend observed with *in situ* hybridization data.

### The rate of protein sequence evolution is associated with ohnolog retention

The nervous system expression bias could be an indirect effect of other factors driving differential retention of duplicate genes. For example, it was observed that genes with slow rates of amino acid substitution were more retained as ohnologs (Davis and Petrov 2004; Brunet et al. 2006). Since genes expressed in the nervous system also tend to be slowly evolving (Duret and Mouchiroud 2000; Gu and Su 2007; Drummond and Wilke 2008; Kryuchkova-Mostacci and Robinson-Rechavi 2015), the rate of amino acid substitutions could be a confounding factor behind the expression bias.

We first verified using our dataset that the 10% genes with the lowest non-synonymous substitutions rate values (*d*_N_, calculated from pairwise comparisons of mouse-rat orthologs) were indeed significantly enriched for expression in nervous structures (Table S12; *p*=8.9e-7, odds=2.13). We also verified that mouse orthologs of zebrafish 3R ohnologs had a lower *d*_N_ than orthologs of singletons (Figure 2A). We then sub-divided the orthologs based on their expression in the nervous system, and surprisingly, while the pattern of slower rate of evolution of orthologs of ohnologs held among nervous system genes, it did not among non-nervous system genes (Figure 2B). This result was confirmed when we split nervous system and non-nervous system genes into 10 equal-sized bins of *d*_N_ (i.e., bins with equal numbers of genes): the proportion of orthologs of ohnologs in each bin was significantly associated with *d*_N_ for nervous system genes, but not so for non-nervous system genes (Figure 3A).

**Figure 2:**
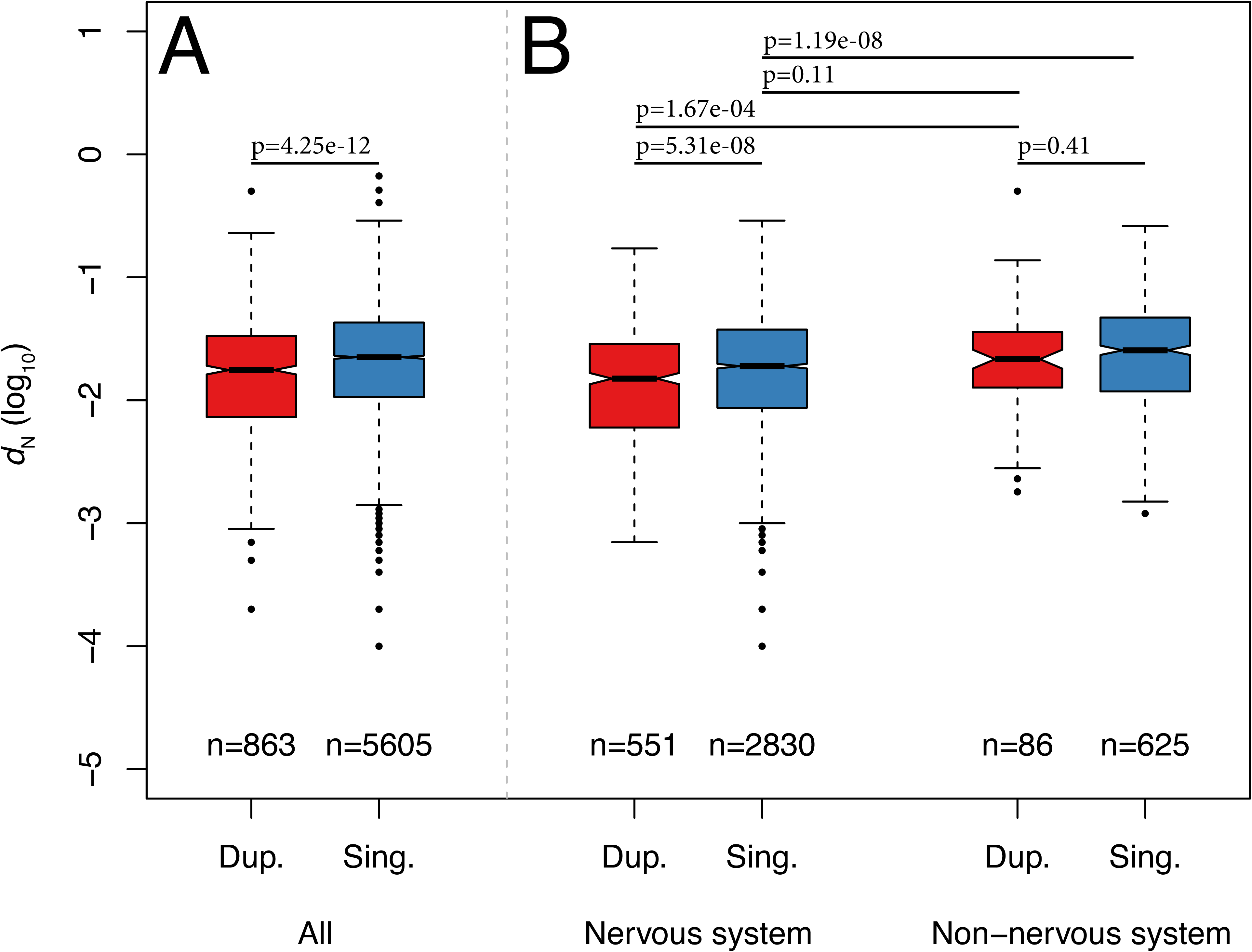
A: Comparison of the rate of protein sequence evolution (*d*_N_, plotted in log_10_ scale) for mouse orthologs of zebrafish 3R ohnologs (“Dup.”) or singletons (“Sing.”). The number of genes in each category is indicated below each box. The *p*-values from a Wilcoxon test comparing categories are reported above boxes. The lower and upper intervals indicated by the dashed lines (“whiskers”) represent 1.5 times the interquartile range, or the maximum (respectively minimum) if no points are beyond 1.5 IQR (default behaviour of R function boxplot). B: similar to (A), but mouse orthologs of zebrafish 3R ohnologs and singletons are split according to their expression in the nervous system (“Nervous system” and “Non-nervous system”). The numbers of duplicates and singletons genes do not add up to numbers of genes in (A) because only genes with *in situ* hybridization data were used for this analysis.

**Figure 3:**
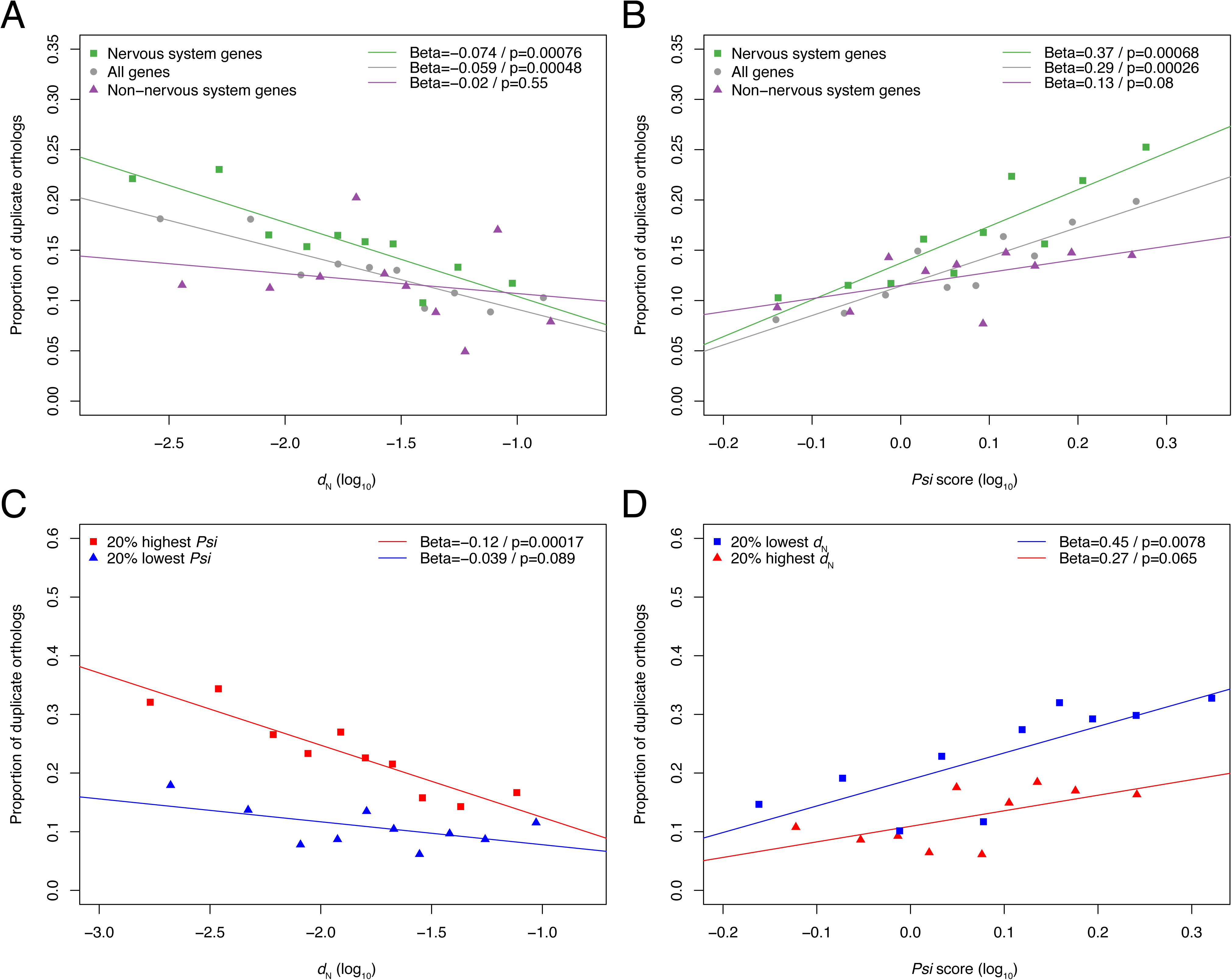
A: Relation between proportion of mouse orthologs of zebrafish 3R ohnologs and rate of non-synonymous substitution. Genes were split into 10 equal-sized bins of *d*_N_, and the median *d*_N_ of each bin was plotted on the x-axis (in log_10_ scale). A linear regression was fit to the 10 data points, whose slope (Beta value) and *p*-value are indicated in the top-right corner of the plot. The analysis including all genes is plotted in grey and circles, while the analysis including only nervous system genes is plotted in green and squares and the analysis including only non-nervous system genes is plotted in purple and triangles. B: Relation between proportion of mouse orthologs of zebrafish 3R ohnologs and Akashi’s test *Psi* score, for nervous and non-nervous system genes. Legend similar to (A). C: Similar to (A), using only nervous system genes, divided in two groups: the 20% genes with lowest *Psi* score plotted in blue and squares, and the 20% genes with highest *Psi* score plotted in red and triangles. D: Similar to (B), using only nervous system genes, divided in two groups: the 20% genes with lowest *d*_N_ plotted in blue and squares, and the 20% genes with highest *d*_N_ plotted in red and triangles.

Since there are more than four times the number of nervous system genes than non-nervous system genes in our analysis, we verified that this results could not be explained by power issues (Supplementary text). We also verified that this pattern held when limiting the set of nervous-system genes to those that were not expressed in any non-nervous structure (Figure S3).

In summary, the biological signal of differences of sequence evolution rates between ohnologs and singletons, apparent at the whole-genome level in our analysis and previously reported in other studies, is in fact mainly caused by nervous system genes. These greatly outnumber non-nervous system genes, for which no sequence evolution rate difference is observed between ohnologs and singletons.

### Highly expressed nervous system genes are more retained as ohnologs

The main hypothesis to explain the slow rates of sequence evolution of nervous-system genes is that their protein sequence was optimized over the course of evolution (Drummond et al. 2005; Drummond and Wilke 2008; Zhang and Yang 2015; Wang 2016). Preferred amino-acids minimize the levels of misfolded proteins, which are toxic to cells because they are prone to aggregate to other proteins and to hydrophobic surfaces such as membranes (Drummond and Wilke 2008; Yang et al. 2012). Preferred amino-acids also minimize the levels of proteins misinteracting with other proteins (Yang et al. 2010).

The long lifetimes and high membrane surface area of neurons make them particularly vulnerable to damages of toxic proteins, explaining why in vertebrates the amino-acid sequences of nervous system genes are the most optimized and conserved (Drummond and Wilke 2008; Drummond and Wilke 2009; Biswas et al. 2016). We verified using our dataset that the negative correlation between *d*_N_ and expression levels was indeed markedly stronger in nervous tissues compared to other tissues (Figure S4A) (Duret and Mouchiroud 2000; Drummond and Wilke 2008; Kryuchkova-Mostacci and Robinson-Rechavi 2015).

We then tested whether the association between *d*_N_ and retention rates could be driven by the expression level of genes in the nervous system. To summarize the expression of genes across nervous system tissues, we considered their mean or maximum expression level across the 102 samples from 46 nervous tissues from the GSE16496 microarray experiment (Figures S4B, S5A and S5B). We then split genes into ten equal-sized bins of mean and maximum nervous system expression level. The proportion of mouse orthologs of zebrafish 3R ohnologs in the bins was significantly associated with maximum, but not with mean nervous system expression (Figure S6A). Interestingly, the association with maximum nervous expression was maintained when we controlled for *d*_N_ by repeating the same analysis on genes with the highest or lowest *d*_N_ (Figure S6B; although the trend for low *d*_N_ genes was slightly below the significance threshold). Conversely the association between *d*_N_ and retention rate disappeared when we controlled for maximum nervous expression by repeating the *d*_N_ analysis on genes with the highest or lowest maximum nervous expression level (Figure S6C). A small residual *d*_N_ trend was visible: if maximum nervous expression perfectly controlled for *d*_N_, the two regression lines would be overlapping on Figure S6B, and they would be flat on Figure S6C.

In summary, for nervous system genes, the maximum level of expression in nervous tissues is clearly associated with higher rates of ohnolog retention. It is difficult though from these results alone to assess whether this association is direct or indirectly caused by the association of both factors to *d*_N_.

### Selective constraints at synonymous sites also influence ohnolog retention

The association of retention rates with *d*_N_, and with maximum nervous expression, suggests a potential link with selection for translational accuracy. In addition to selection for optimal amino-acid sequences of genes, which reduces the levels of toxicity of protein products, synonymous codon usage was also shown to be optimized to increase translational robustness and reduce the levels of mistranslation-induced protein products toxicity (Drummond and Wilke 2008; Yang et al. 2010). Codons binding their cognate tRNA with higher affinity than non-cognate competitors are translated more accurately, decreasing the chances of incorporation of wrong amino acids during translation. The selection to maintain a state of optimized synonymous codon usage is apparent in the association between *d*_N_ and *d*_S_ (Figure S5C), and in the stronger negative correlation between the rate of synonymous substitutions (*d*_S_) and expression levels in nervous tissues compared to other tissues (Figure S7A). However we could not observe any relation between the proportion of mouse orthologs of zebrafish 3R ohnologs and *d*_S_ (Figure S7B).

This may be explained by the fact that selection for translational robustness does not act on synonymous codon usage at all sites, but predominantly on those that are the most important for the structure of the protein (Drummond and Wilke 2008; Zhou et al. 2009) or that are aggregation-prone (Lee et al. 2010). The strength of this effect can be quantified using Akashi’s test, assessing how strong is the association between preferred codons and conserved amino acids, taken as a proxy of constrained sites in the protein (Akashi 1994).

The odds ratio reflecting this association (“*Psi*” score) was only weakly correlated with both the rate of non-synonymous substitutions, *d*_N_, and with maximum nervous expression (Figure S5E and F), but was more strongly associated with translation rates, calculated from ribosome profiling data in embryonic stem cells, embryonic fibroblasts or neutrophils (Dana and Tuller 2014)(see Materials and Methods; Figure S8). Genes showing the 10% highest *Psi* score were enriched for expression in the nervous system (*p*=0.00081, odds=1.98), and interestingly the top structures were almost exclusively developing ectodermal or neural structures (e.g., “rhombomere”, “presumptive midbrain”, “limb bud”; Table S13). In summary, the *Psi* score captures some aspect of selection for translational accuracy that seems largely independent of the constraints on amino acid sequences.

Interestingly, when we separated mouse genes in 10 equal-size bins of *Psi*, we observed a significant relation with the proportion of orthologs of ohnologs (Figure 3B). And similarly to the *d*_N_ trend, the association was supported for nervous system genes, but not for non-nervous system genes. Focusing on the nervous system genes, we checked whether the *d*_N_ and *Psi* trends were dependent (Figures 3C and D). The proportion of orthologs of ohnologs was the highest for the genes with the lowest *d*_N_ and the highest *Psi*, suggesting a positive interaction between the effects of *d*_N_ and *Psi*. Nervous system genes that had either high *d*_N_ or low *Psi* included around 10% of orthologs of ohnologs, similarly to non-nervous system genes, but nervous system genes that had both low *d*_N_ and high *Psi* included more than 30%.

In summary, genes with the most optimized sequences, both at non-synonymous and at synonymous sites, were more retained in duplicate after whole-genome duplication.

### The nervous system bias is independent from the dosage-balance hypothesis

The dosage-balance hypothesis was previously proposed to explain ohnolog retention after whole-genome duplication (Freeling and Thomas 2006; Makino and McLysaght 2010; Birchler and Veitia 2012; Singh et al. 2012). Groups of interacting genes, sensitive to relative dosage changes (e.g., members of a protein complex, or genes belonging to the same metabolic pathway) could be maintained in duplicates because the loss of any gene of the group would lead to dosage imbalance and be detrimental (Birchler and Newton 1981; Birchler et al. 2005). Notably, dosage imbalance is expected to impact the formation of protein complexes involving at least two different genes, and composed of at least three subunits (Veitia et al. 2008), so genes involved in such complexes should be more retained after whole-genome duplication.

We tested this using protein complex data from the UniProtKB/Swiss-Prot database (The UniProt Consortium 2015), where complex type and number of subunits are precisely annotated. We split genes into six groups: genes involved in (i) monomers, (ii) homomultimers and (iii) hetero-dimers, which should not be sensitive to dosage imbalance; (iv) genes involved in hetero-multimers with at least 3 subunits, which should be sensitive to dosage imbalance; (v) genes involved in uncharacterized complexes, which likely include some genes sensitive to dosage imbalance; and finally (vi) non-annotated genes. Genes annotated to several groups were kept in the group expected to be most sensitive to dosage imbalance (see Materials and Methods). We observed that the proportion of mouse orthologs of zebrafish 3R ohnologs was the highest for members of hetero-multimer complexes, consistent with the dosage-balance hypothesis (Figure S9A). This effect was independent from nervous system expression, since it was observed both for nervous and non-nervous system genes. When controlling for *d*_N_, we observed a positive interaction between the two effects: members of hetero-multimer complexes that had low *d*_N_ were the most retained (Figure S9B).

Another manifestation of the dosage-balance hypothesis could be selection to maintain stoichiometry within metabolic pathways. Previous studies in *Paramecium* and *Arabidopsis* have reported that the retention rate of genes involved in metabolic pathways differed across timescales, and was higher than other genes for recent whole-genome duplication events, while it was lower for ancient events (Aury et al. 2006; Gout et al. 2009; Bekaert et al. 2011). Consistent with these observations, we observed a lower proportion of mouse orthologs of zebrafish 3R ohnologs among genes involved in metabolic processes (Figure S10), both for nervous and non-nervous system genes, confirming that there is no long-term action of selection against dosage imbalance on whole pathways.

Finally, we examined the relation between the level of protein connectivity and retention rates. The number of protein-protein interactions was previously taken as a proxy for sensitivity to dosage imbalance (Prachumwat and Li 2006; Flagel and Wendel 2009; Rodgers-Melnick et al. 2013; Cuypers and Hogeweg 2014). We observed that mouse orthologs of zebrafish 3R ohnologs had a significantly higher connectivity than orthologs of singletons (Figure S11A), in agreement with previous studies (Hakes et al. 2007; Liang and Li 2007; Rodgers-Melnick et al. 2012). But similarly to the *d*_N_ trend, when we sub-divided genes based on their expression pattern, the trend held only for nervous system genes. Since highly connected genes tend to display a lower *d*_N_ (Figure S5G), we tested whether the relation between retention rate and connectivity could be explained by *d*_N_ differences: this was not the case, with connectivity and *d*_N_ even positively interacting to explain retention rates (Figure S11B and C). Similarly, the connectivity trend could not be explained by other weakly correlated factors, maximum nervous expression and *Psi* score (Figure S5I and H), but it disappeared when we split genes based on their annotation the six complex subtypes (Figure S11D).

This suggests that the connectivity trend could indeed be explained by higher dosage sensitivity of most highly connected genes, although there is no clear *a priori* reason for the trend to be seen only among nervous system genes (see Discussion).

### Small-scale duplication is not associated to the same underlying factors

Small-scale duplicates have often been observed to behave in an opposite way to ohnologs, a pattern that we confirmed with the lower rate of duplication of genes expressed in the nervous system (Fig. 1B). More careful examination indicated that this bias was not caused by the same underlying factors as ohnologs.

First, there was a difference in *d*_N_ values between genes that experienced a small-scale duplication event and other genes, but this was true both for nervous system and non-nervous system genes (Figure S12A). This suggests that the depletion for nervous system expression might just be an indirect consequence of the association between small-scale duplication and sequence evolution rates. Second, the relation between the proportion of small-scale duplicates and *d*_N_ was best explained by a linear fit, whereas the best model was a log-linear trend for ohnologs (Figure S12B). Third, there was no relation between the proportion of small-scale duplicates and *Psi* score, suggesting that there was no association between selection for translational accuracy and small-scale duplication patterns (Figure S12B and C).

Since expression level is a major determinant of *d*_N_, we examined the relation between the proportion of small-scale duplicates and summaries of expression levels across nervous and non-nervous tissues. The best trend was obtained using the average expression level across all available tissues (not only nervous tissues; Figure S13A). There was even a positive interaction between the effects of *d*_N_ and average expression level: genes with a small-scale duplication history displayed both a low average expression and a high *d*_N_ (Figure S13B and C).

We finally tested the dosage-balance hypothesis on small-scale duplicates, using protein complex information. In comparison to whole-genome duplication patterns (Figure S9), hetero-multimer genes displayed a lower proportion of duplicates, consistent with their dosage-sensitivity (Figure S14), but this was only true for nervous system genes. For the subset of non-nervous system genes, the proportion of duplicate hetero-multimer genes was surprisingly higher than the other groups of genes. Given that this category includes the lowest number of genes, this pattern must be interpreted carefully. The low number of small-scale duplicates makes it unfortunately difficult to reliably test the dependency of this trend with respect to the *d*_N_ and average expression level trends.

## Discussion

In this study we took advantage of thousands of high quality *in situ* hybridization data describing precisely mouse and zebrafish gene expression patterns. These are mapped to ontologies describing the anatomy of these species, making it possible to perform ontology enrichment tests and to detect tissues enriched for the expression of genes of interest. This methodology corrects for biases in annotation and in data availability, i.e., some anatomical structures are better annotated than others (Yon Rhee et al. 2008).

We uncover a strong and robust trend whereby genes expressed in neural tissues are more likely retained in duplicate after whole-genome duplication. These same genes are less likely to duplicate via small-scale duplication events. To our knowledge, this result was never previously reported, but is fully consistent with previous studies. For example, ohnologs were found enriched for Gene Ontology terms related to signaling, behavior, neural activity or neurodevelopment (Brunet et al. 2006; Putnam et al. 2008; Roux and Robinson-Rechavi 2008; Kassahn et al. 2009; Huminiecki and Heldin 2010; Howe et al. 2013; Schartl et al. 2013), which are typical nervous system genes functions. The slow rate of sequence evolution of ohnologs (Davis and Petrov 2004) can also be explained by the tendency of nervous system genes to evolve slowly (Duret and Mouchiroud 2000; Gu and Su 2007; Drummond and Wilke 2008).

Surprisingly, there have been few previous analyses of ohnolog retention biases with respect to gene expression patterns, probably because of the limited anatomical resolution of most microarray and RNA-seq datasets, and the difficulty in gathering many *in situ* hybridization experiments for an integrated analysis. Satake and colleagues (2012) reported that the proportion of 2R ohnologs detected in EST datasets was the highest in ectoderm-derived tissues, while the proportion of small-scale duplicates was the lowest, which is consistent with our observations.

Once this pattern was established, the next challenging task was to disentangle, within the network of factors associated with retention rates, which factors could be causal, and more broadly, which mechanisms are in action (Drummond et al. 2006; Singh et al. 2012; Kryuchkova-Mostacci and Robinson-Rechavi 2015). These analyses were done in mouse, an outgroup species used as proxy for the pre-duplication state in the teleost fish ancestor. Unfortunately we lack at present good data to verify these patterns in teleosts, i.e. we lack closely related genomes to zebrafish, or at least fish genomes outgroup to the 3R whole-genome duplication with sufficient functional data.

The rate of non-synonymous substitutions, *d*_N_, is strongly associated to the maximum level of expression across nervous tissues, an association that is likely caused by selection for optimized amino-acid sequences against the toxic effects of misfolded or misinteracting protein products (Drummond and Wilke 2008; Zhang and Yang 2015). Interestingly, both factors are independently associated with retention of nervous system ohnologs, suggesting that selection for optimized amino-acid sequences could play a key role in this process. This hypothesis is corroborated by the observation that another manifestation of selection against toxic protein products, the optimization of codon usage at structurally sensitive sites to increase translational robustness, is also associated with retention rates, and this effect is not controlled by *d*_N_ or maximum nervous system expression. All these effects even seem to be positively interacting: genes that have a low *d*_N_, a high maximum nervous system expression and a high *Psi* score have the highest chances of retention.

After whole-genome duplication, duplicate gene loss starts with the fixation of loss-of-function mutations in one of the gene copies. This can occur neutrally as long as the gene function is backed-up by the other copy. Thereafter the non-functional copy accumulates other substitutions and degenerates (Albalat and Canestro 2016). Such a neutral scenario might not be possible for nervous system genes whose sequence is constrained by selection. Indeed, mutations occurring both at non-synonymous sites, and at some synonymous sites, can increase the rate of production of toxic proteins, and this deleterious effect should hamper their fixation in the population (Figure 4). This simple model can explain how both duplicate gene copies can be “protected” from degeneration by purifying selection after whole-genome duplication, despite functional redundancy.

**Figure 4:**
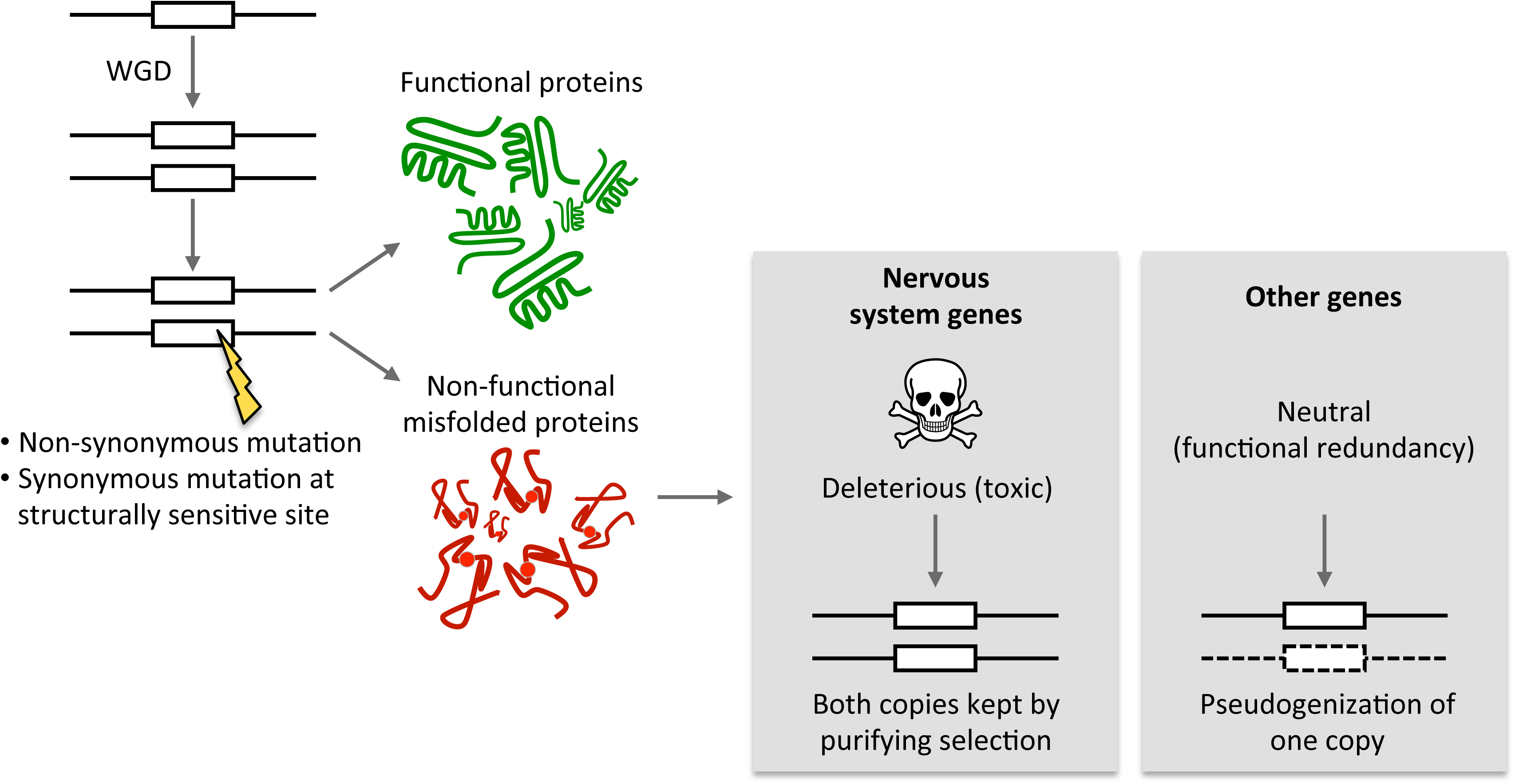
Illustration of the model proposed to explain favored retention of nervous system genes after whole-genome duplication. Non-synonymous mutations or synonymous mutations at structurally sensitive sites on one duplicate copy can cause an increase in the production of non-functional toxic protein products. This is likely neutral in most tissues, since the function loss is backed-up by the other copy. This could however be deleterious for nervous system genes because they are expressed in non-renewing cells, sensitive to the toxic effects of misfolded or misinteracting proteins. Purifying selection will thus prevent the fixation of such mutations, and indirectly contribute to the preservation of both ohnologs.

More broadly than nervous system genes, any gene subject to dominant deleterious effects mutations should be more likely retained as ohnolog, since the organism would pass by a low fitness intermediate when losing one copy. In fact, Gibson and Spring (1998), and later Singh and colleagues (2012) proposed such a model to explain the puzzling observation that disease-causing genes were preferentially retained after the 2R whole-genome duplications (Gibson and Spring 1998; Makino and McLysaght 2010; Dickerson and Robertson 2012; Singh et al. 2012; Chen et al. 2013; Malaguti et al. 2014; Singh et al. 2014; Tinti et al. 2014). They later supported this model by theoretical population genetics work (Malaguti et al. 2014), explaining the accumulation of repertoires of “dangerous” genes after whole-genome duplication. This is also supported by the enrichment of ohnologs among genes for which copy number variants are pathogenic (Rice and McLysaght 2017).

Of course, our model does not totally exclude the possibility of pseudogenization of nervous system genes. For example, a mutation introducing a stop codon at the very beginning of the coding sequence is not likely to produce toxic products. It is also possible that regulatory mutations first silence one duplicate copy, opening the way to its neutral degeneration (Thompson et al. 2016). The neutral evolution of asymmetric expression levels between duplicate copies has indeed been reported (Gout and Lynch 2015; Lan and Pritchard 2016; Thompson et al. 2016). But (i) this process was shown to require substantial amounts of time, and (ii) the evolution of expression levels in the nervous system is tightly controlled and slower than in other tissues (Pennacchio et al. 2006; Brawand et al. 2011; Barbosa-Morais et al. 2012; Merkin et al. 2012). Hence, there is little reason to think that other pseudogenization routes would compensate the deficit of losses for nervous system ohnologs.

Our model also does not exclude the possibility that some ohnolog pairs are retained through the action of previously described mechanisms (Innan and Kondrashov 2010), for example sub-functionalization (Force et al. 1999) or neo-functionalization (Ohno 1970; He and Zhang 2005). We could not find any reasonable explanation for the nervous system retention bias using these alternative mechanisms, but these might however be necessary to maintain ohnologs in the long term. For example, nervous system duplicates that avoided rapid initial loss could be eventually retained because they evolved new functions later in time (Kassahn et al. 2009; Chen et al. 2011).

Another interesting model is the dosage-balance hypothesis, which was proposed to be a major determinant of duplicate gene retention after whole-genome duplication, at least on short evolutionary time scales (Papp et al. 2003; Freeling and Thomas 2006; Makino and McLysaght 2010; McLysaght et al. 2014; Thompson et al. 2016). This hypothesis is difficult to test in vertebrates because there are only a few noisy datasets allowing to assess the sensitivity of genes to dosage imbalance. Previous studies have sometimes relied on indirect evidence; for example, it was found that genes with high levels of protein-protein interactions (more connected genes) tended to be more retained after whole-genome duplication, which was interpreted as an evidence that these genes are more sensitive to changes in dosage (Liang and Li 2007; Rodgers-Melnick et al. 2013).

Such an interpretation is subject to caution. Although we indeed found connectivity to be significantly associated with retention rates, we noticed that the trend was only supported for nervous system genes. There is *a priori* no reason to expect this pattern from the dosage-balance hypothesis. However, it was shown that amino-acid sequences are optimized to reduce the levels of misinteraction with other proteins (Yang et al. 2012), an effect that might be more important for highly connected proteins, and for those expressed in non-renewing neural cells than other cell types. Protein surface residues in particular are optimized for decreased stickiness and misinteractions, which are deleterious because they waste functional molecules, can interfere with functional interactions, or initiate damaging cellular processes (Zhang et al. 2008; Vavouri et al. 2009; Yang et al. 2012). The chances of detrimental effect might be higher for highly connected proteins, and similarly to protein misfolding, the effects might be more detrimental to non-renewing neural cells than other cell types, contributing to a retention bias of highly connected nervous system ohnologs. Hence, this mechanism provides an alternative explanation, probably complementary to the dosage-balance hypothesis, to the relation between connectivity and retention rates.

A better source of evidence to test the dosage-balance hypothesis is protein complex data. But different complex subtypes are not equally sensitive to dosage imbalance (Veitia et al. 2008). When separating complexes into permanent or transient complexes, a previous study in human surprisingly observed that the retention rates after the 2R whole-genome duplications were lower for permanent complexes, despite their higher susceptibility to dosage-balance constraints (Singh et al. 2012). We separated genes into those involved or not in dosage-sensitive complexes and, consistent with the dosage-balance hypothesis, observed a higher retention of the former. Moreover, this trend was supported both for nervous system and non-nervous system genes, and was independent of confounding factors such as *d*_N_, suggesting that the effects of selection against gene dosage imbalance on ohnologs retention are likely independent from the effects of selection against toxic protein products.

Another analysis that we performed with another source of data gave somewhat conflicting results, but the annotations were less comprehensive and precise (see Supplementary Text). This underlines that careful analysis and high-quality datasets are needed to study the effects of selection against gene dosage imbalance on ohnologs retention independently from the effects of selection against toxic protein products. For example, it is important to be careful with data transferred across species, which could be biased by the rates of sequence evolution, and to study pre-duplication biases in an outgroup species, because duplication itself likely influences post-duplication evolution of dosage-sensitive genes (e.g., ohnologs might be more likely to evolve into hetero-multimer complexes members)(Musso et al. 2007; Qian and Zhang 2014).

Interestingly, it is worth noting that the dosage-balance hypothesis, quite similarly to our model of Figure 4, also explains duplicate retention biases by the action of purifying selection (Freeling and Thomas 2006), acting not on detrimental mutations in coding sequences as in our model, but also on detrimental changes in expression of dosage-sensitive genes. The predominant role of purifying selection can account for the observation that ohnologs usually do not duplicate via small-scale duplication events. Indeed, small-scale duplication events first need to reach fixation in the population, a process that is rarely successful for such genes, whose mutations can be dominant negative (Innan and Kondrashov 2010; Singh et al. 2012).

A recent study (Rice and McLysaght 2017) has reported that genes found in pathogenic copy number variant mutations are involved in development, enriched in protein complexes, have high expression, and have evolutionary patterns depleted in small-scale duplications but enriched in ohnologs. These observations are consistent with dosage-balance, and interpreted in that manner (Rice and McLysaght 2017). Yet, interestingly, the “class P” (pathogenic) genes of Rice and McLysaght (2017) have expression highly enriched in nervous system structures by TopAnat (not shown). It is possible that dosage imbalance effects might be more severe in the nervous system than other tissues, but (i) there is to our knowledge no prior report of this effect, and (ii) we did not observe an under-representation of small-scale duplications in nervous system ohnologs compared to non-nervous system ohnologs (Supplementary Text). Thus the observations of Rice and McLysaght (2017) could rather be at least in part explained by our hypothesis of selection against the toxicity of protein products.

We observed that small-scale duplicates were rarely expressed in the nervous system, but this time, likely as an indirect effect of low fixation and retention rates of duplicates of slowly evolving highly expressed genes. This is consistent with purifying selection acting primarily on the deleterious effects of doubling the gene expression induced by small-scale duplications (Schuster-Böckler et al. 2010; McLysaght et al. 2014; Rice and McLysaght 2017). Although average expression level is highly correlated with *d*_N_, it did not account for the entirety of the relation between *d*_N_ and rate of small-scale duplication. The additional effect of *d*_N_ could be due to post-duplication biases, that we did not control for in this analysis. Small-scale duplicates were indeed shown to experience an accelerated evolutionary rate after duplication, possibly associated with a process of sub- or neo-functionalization (Jordan et al. 2004; Fares et al. 2013; Pegueroles et al. 2013). Finally, selection against protein misfolding was not associated with small-scale duplication rates. This is perhaps not surprising, because the sequence of genes expressed in tissues sensitive to protein misfolding was optimized by natural selection, and duplication is unlikely to affect this, especially since the fixation phase of duplicates is probably too short for point mutations to accumulate.

## Conclusion

The implications of our results are manifold. First, they confirm that whole-genome duplication is a unique type of evolutionary event, which enriches the gene set of a lineage with genes under strong purifying selection, e.g., dosage-sensitive genes, disease-causing genes, or nervous system genes. Mutations affecting the sequences or the expression of these genes can have clear detrimental consequences, adding a long term burden to genomes. Counter-intuitively, the preferential retention of these genes is driven by the action of purifying selection alone, although this is usually viewed as a protective force. Our study focused on vertebrates, but such a situation is most likely true for other organisms which experienced whole-genome duplications, such as plants or unicellular eukaryotes, although the sets of retained genes might differ.

On the other hand, whole-genome duplications have often been claimed to be beneficial in the long term, since the addition of new genes to genomes provides new material for evolution to act on, and increases evolvability of the lineages (Van de Peer et al. 2009; Kondrashov 2012; Cuypers and Hogeweg 2014). A particularly interesting example is the ancestral 2R event, which added to the genomes of vertebrates a large number of regulatory genes, such as transcription factors, as an indirect effect of purifying selection for gene dosage balance. Freeling and Thomas coined this phenomenon a “spandrel” of purifying selection, and suggested that it contributed to the increased morphological complexity of vertebrates (Gould and Lewontin 1979; Freeling and Thomas 2006).

Our results highlight that another such by-product of purifying selection is the enrichment of the vertebrate genomes for nervous system genes, at a time which coincided with major evolutionary novelties of the nervous system. The expanded toolkit of nervous genes likely provided opportunities for regulatory network rewiring and new functions to evolve (Evlampiev and Isambert 2007; Oakley and Rivera 2008; Chakraborty and Jarvis 2015). For example, it was suggested that the 2R events gave vertebrates the tools to evolve new structures such as the neural crest, placodes and a midbrain–hindbrain boundary organizer (Holland 2009). Similarly, in fish it was suggested that the 3R whole-genome duplication contributed to expand the toolkit of cognition-related genes that gave teleosts a high level of behavioral complexity compared to other groups of cold-blooded vertebrates such as amphibians and reptiles (Schartl et al. 2013).

## Materials and Methods

Data files and analysis scripts are available on our GitHub repository: <u>http://github.com/julien-roux/Roux_Liu_and_Robinson-Rechavi_2016</u>

### Mouse and zebrafish *in-situ* hybridization data

Mouse (*Mus musculus*) RNA *in-situ* hybridization expression data were retrieved from the GXD database (Smith et al. 2007; Smith et al. 2014) in December 2014. Wild-type data, obtained under non pathological conditions, and with no treatment (“normal” gene expression) were integrated into Bgee (http://bgee.org/), a database allowing the comparison of transcriptome data between species (Bastian et al. 2008). The data used in this article all come from the release 13 of Bgee. In Bgee, expression data are mapped to the Uberon anatomical ontology (http://uberon.org). The mapping from the EMAP (Bard et al. 1998) and MA (Hayamizu et al. 2005) mouse anatomical ontologies (onto which GXD *in-situ* hybridization data are mapped) to Uberon was obtained from Uberon cross-references. Terms from the EMAPA and MA ontologies that were not present in the Uberon ontology, but to which *in-situ* hybridization data were mapped were also included in the analyses.

Similarly, zebrafish (*Danio rerio*) *in-situ* hybridization expression data were retrieved from the ZFIN database (Sprague et al. 2006) in December 2014 and integrated into Bgee release 13 after mapping to the Uberon anatomical ontology. Terms from the ZFA ontology that were not present in the Uberon ontology, but to which *in-situ* hybridization data were mapped were also included in the analyses.

### Mouse microarray data

Mouse microarray data and their mapping to the Uberon anatomical ontology were retrieved from Bgee release 13. We targeted experiments including a large number of samples from many different tissues, and including multiple nervous and non-nervous system tissues. We retained the accessions GSE3594, GSE10246 and GSE16496.

GSE3594 is a dataset composed of 129 samples from 24 neural tissues and 10 body tissues from different strains of inbred mice (Zapala et al. 2005). This experiment was hybridized to the Affymetrix Murine Genome U74A Version 2 array. Raw data (CEL files) were not available from GEO, so the normalized intensities and present/absent calls provided by the MAS5 software (Liu et al. 2002) were used.

GSE10246 corresponds to the GNF Mouse GeneAtlas V3 (Su et al. 2004) and there were 91 samples from 45 tissues (including 12 neural tissues, as well as 7 sub-structures of the eye) included into Bgee. This dataset was hybridized to the Affymetrix Mouse Genome 430 2.0 Array chip and was reprocessed through the Bgee pipeline (see http://bgee.org/bgee/bgee?page=documentation). Briefly, this includes normalization of the signal of the probe sets by the gcRMA algorithm, and a Wilcoxon test on the signal of the probes ets against a subset of weakly expressed probe sets to generate present/absent calls (Schuster et al. 2007).

GSE16496 included expression data 102 samples from 46 regions of the mouse central nervous system (Kasukawa et al. 2011). This dataset was hybridized to the Affymetrix Mouse Genome 430 2.0 Array chip and also reprocessed through the Bgee pipeline.

We summarized the expression of genes across nervous system tissues by considering for each gene the mean, median or maximum of log_2_ signal across all samples from the GSE16496 experiment. Results were similar when using nervous tissue samples of the GSE3594 (not shown). Because results were similar using the median or the mean expression across nervous tissues, we only show results using the median.

### Human RNA-seq data

Human RNA-seq data from the GTEx consortium (Melé et al. 2015; The GTEx Consortium 2015) were retrieved from Bgee release 14 (GTEx processed and annotated data available on ftp.bgee.org; full release planned in Feburary 2017). All samples were manually annotated to the Uberon ontology and only healthy samples were retained, based on metadata annotation (e.g., medical history or cause of death). There were 4,860 retained GTEx libraries, mapped to 75 different Uberon terms. The libraries were reprocessed through the Bgee pipeline to generate present/absent calls for each gene. Briefly, RNA-seq reads from each library were pseudo-aligned with Kallisto (version 0.42.4)(Bray et al. 2016) to the annotated human transcripts from Ensembl (release 84). Transcript-level TPM estimates were then summed at the gene level. Reads were also pseudo-mapped to a set of 28,573 intergenic regions, located at least 500 bp away from any genic region, and whose size ranged from 2000 bp to 20,000 bp. The “background” expression signal observed at these regions was used to set a TPM threshold for each library to determine presence/absence calls. At the threshold the ratio of the proportions of intergenic regions called present to the proportion of coding genes called present was set to 5%.

### Identification of duplicates and singletons

Gene families were obtained from the Ensembl database release 79 (Hubbard et al. 2009). We used the Perl API to query the Ensembl Compara Gene trees (Vilella et al. 2009) and scan for gene trees with specific topologies. Notably we stringently selected sets of genes with or without duplications on specific branches of the vertebrate phylogenetic tree. We randomly picked a subset of gene trees to verify that they indeed displayed the expected topologies. Below is a description of the selected topologies, which are illustrated on Figure S1. These are dependent on the set of species integrated into Ensembl release 79, accessible at http://mar2015.archive.ensembl.org/info/about/speciestree.html. All genes lists are available as supplementary material (file gene_lists.zip) and scripts are available on our GitHub repository.

### Fish-specific (3R) whole-genome duplication (Figures S1A, B, C and D)

We first selected subtrees with a basal speciation node dated at the Neopterygii taxonomical level. These subtrees include a spotted gar (*Lepisosteus oculatus*) outgroup, which did not experience the 3R duplication (Braasch et al. 2016), and teleost fish species, which experienced it. We classified zebrafish genes as confident 3R duplicates if the child node of the root of the subtree was a high confidence (score above 50%) duplication node dated at the Clupeocephala taxonomic level, followed by two speciation nodes dated at the Clupeocephala taxonomic level, each delineating a subtree containing no further duplication or loss on the branches leading to zebrafish (i.e., one zebrafish gene per subtree). We classified zebrafish genes as confident 3R singletons if the child node of the root of the subtree was a speciation node dated at the Clupeocephala taxonomic level, with no further duplication or loss on the branches leading to zebrafish. In total we obtained 2422 ohnologs, and 8973 singletons.

Of note, our identification of ohnologs is based on phylogeny alone, and does not use any synteny information. Small-scale duplicates that emerged on the Clupeocephala branch will be wrongly incorporated in the list of 3R ohnologs. Given relatively low rate of retention of duplicates originating from small-scale duplication (Lynch and Conery 2000), we ignored this problem in our analyses.

We classified mouse or human genes as confident orthologs of zebrafish 3R ohnologs if there was a two-to-one orthology relationship to a single mouse/human gene at the Euteleostomi taxonomical level. We classified mouse or human genes as confident orthologs of zebrafish 3R singletons orthologs if there was a one-to-one orthology relationship to a single mouse/human gene at the Euteleostomi taxonomical level. In total we obtained 974 mouse orthologs of 3R ohnologs, 6,373 mouse orthologs of 3R singletons, 976 human orthologs of 3R ohnologs, and 6,358 human orthologs of 3R singletons.

### Vertebrate (2R) whole-genome duplications (Figures S1E and F)

It is still debated whether one or two whole-genome duplication events occurred at the base of vertebrates (Smith and Keinath 2015). In gene trees, we thus allowed for the possibility of one or two duplications at the base of vertebrates. If two rounds of wholegenome duplication really occurred, this means that we required ohnologs of at least one event to be retained.

We first selected subtrees with a basal speciation node dated at the Chordata taxonomical level – or at the Bilateria taxonomical level when there was no chordate node in the subtree. We classified mouse genes as confident 2R duplicates if the child node of the root of the subtree was a high confidence duplication node dated at the Vertebrata taxonomic level, followed by an optional second high confidence duplication node dated at the Vertebrata taxonomic level, followed by two speciation nodes dated at the Vertebrata taxonomic level, each delineating a subtree containing only one mouse gene and including genes from at least two different fish species. We used Euteleostomi instead of Vertebrata to date the 2R duplications if there was no lamprey gene in the subtree. We classified mouse genes as confident 2R singletons if the child node of the root of the subtree was a speciation node dated at the Vertebrata/Euteleostomi taxonomic level, and delineated a single subtree including one mouse gene and genes from at least two different fish species. We could not enforce strictly the constraint that no duplication occurred in the tetrapod lineage on the branches leading to mouse, because Ensembl mammalian trees include a high number of dubious duplication nodes (duplication confidence score = 0) that are generated when the gene tree topology is not consistent with the species tree. Given the high number of mammalian species in Ensembl, this problem occurred in virtually each of the trees we examined. In total, we obtained 1389 2R ohnologs and 2999 singletons.

### Small-scale duplications (Figures S1G and H)

We observed that genome assembly and annotation errors resulted in a high number of likely artifactual species-specific paralogs in gene trees. Thus we chose to retain only small-scale duplicates that originated before the split with at least one species. For zebrafish the more recently diverged sister species present in Ensembl was the cave fish *Astyanax mexicanus*, so we focused on small-scale duplicates that originated on the Otophysa branch (deeper branches could not be considered because of the 3R fish-specific genome duplication). We first selected subtrees with a basal speciation node dated at the Clupeocephala taxonomical level. We then retrieved homology relationships between all zebrafish paralogous genes in the subtree (if any), and retained only the high-confidence ones, which did not involve paralogs with 100% sequence identity (probable assembly artifacts) or <10% sequence identity (probable gene split), and were dated at the Otophysa taxonomical level. In total we obtained 385 duplicates.

For mouse we focused on mammal-specific small-scale duplications. We first selected subtrees with a basal speciation node dated at the Mammalia taxonomical level. We then retrieved homology relationships between all mouse paralogous genes in the subtree (if any), and retained only the high-confidence ones, which did not involve paralogs with 100% sequence identity or <10% sequence identity, and were dated at the Theria, Eutheria, Boreoeutheria, Euarchontoglires, Glires, Rodentia, Sciurognathi, or Murinae taxonomical levels. In total, we obtained 646 duplicates.

### Ontology enrichment analyses

Enrichment and depletion of expression in anatomical structures were tested with a Fisher exact test using a modified version of the R Bioconductor package topGO (http://bioconductor.org/; Adrian Alexa, pers. comm.)(Gentleman et al. 2004; Alexa et al. 2006; R Development Core Team 2007), allowing to handle other ontologies than the Gene Ontology. We defined the reference set as all the genes for which we had expression data in at least one structure of the organism across all life stages using *in-situ* hybridization data. This accounted for 9398 genes in zebrafish and 11322 genes in mouse, expressed in respectively 1067 and 2783 anatomical structures. Only anatomical structures with annotated expression of at least 5 genes were analyzed.

The expression data were propagated to parent structures in the ontology (e.g., a gene expressed in the “hindbrain” was also considered expressed in the parent structure “brain”), a methodology that is very helpful to automatically integrate large amounts of implicit knowledge. However this can result in the enrichment of non-independent terms, and of top-level terms of the ontology that are sometimes difficult to interpret, a behavior that is well known for Gene Ontology enrichment tests (Alexa et al. 2006; Falcon and Gentleman 2007; Yon Rhee et al. 2008). To correct for this effect, we used the “weight” algorithm available in the topGO package, a bottom-up approach that up or down-weights terms depending on whether they benefit from the signal of their children structures (Alexa et al. 2006). Unless explicitly mentioned, this algorithm was used in the paper. Using another decorrelation algorithm of the topGO package, the “elim” algorithm, gave similar results (not shown).

A False Discovery Rate correction was applied on the list of *p*-values from tests on all anatomical structures. Structures enriched or depleted with a FDR < 10% are reported (Benjamini and Hochberg 1995). Of note, all analyses in this paper are reproducible using the TopAnat webservice available at http://bgee.org/?page=top_anat#/, as well as programmatically, using the BgeeDB Bioconductor package available at http://bioconductor.org/packages/release/bioc/html/BgeeDB.html. An example script is available as Supplementary material and on our GitHub repository (file expression_enrichment_with_BgeeDB.R). The results from the webservice and the Bioconductor package can differ slightly from our results due to slight differences on the handling of anatomical ontologies.

### List of nervous system anatomical structures

A reference list of anatomical structures belonging to the nervous system in zebrafish and mouse was extracted from the Uberon ontology (as used in the Bgee database release 13). Because it was sometimes debatable whether a structure belonged to nervous system or not (e.g. sensory organs), we created a “strict” list and a “broad” list.

In zebrafish, the strict list included the “nervous system” structure (UBERON:0001016), as well as its sub-structures in the ontology. The “sensory system” structure (UBERON:0001032) and its sub-structures were removed. The broad list included them, as well as presumptive neural structures during development and their sub-structures (future nervous system, UBERON:0016880; neurectoderm, UBERON:0002346) and the structure “neurovascular bundle” and its sub-structures (UBERON:0016630).

In mouse, we used the same criteria, but we also noticed that some structures added to Uberon from the mouse-specific ontologies (EMAPA and MA ontologies) were not connected to any nervous system Uberon term at time of study. We thus added the following list of structures and their sub-structures to our broad list: nerves of urethra (EMAPA:31569), head or neck nerve or ganglion (MA:0000572 and MA:0000580), nerve of prostatic urethra (EMAPA:32279), nerves of urogenital sinus (EMAPA:31533), tail nervous system (EMAPA:16753), testicular branch of genital nerve (EMAPA:29731), nerve of prostate gland (EMAPA:32285), renal cortical nerves (EMAPA:31319), renal medullary nerves (EMAPA:31354), nerve of bladder (EMAPA:31526), nerve of pelvic urethra (EMAPA:31558), and nerve of caudal urethra (EMAPA:31557). Note that many of these species-specific structures are connected to Uberon nervous system structures in the most recent release of Uberon.

The reference lists of nervous system structures were intersected with the list of anatomical structures showing expression of at least 5 genes, to keep only structures for which expression enrichment was effectively tested.

### List of anatomical structures from other systems

We selected the high-level terms in the ontologies corresponding to these broad anatomical systems on zebrafish and mouse: Biliary system (UBERON:0002294), Circulatory system (UBERON:0001009), Digestive system (UBERON:0001007), Exocrine system (UBERON:0002330), Hematopoietic system (UBERON:0002390), Immune system (UBERON:0002405), Musculoskeletal (system UBERON:0002204), Renal system (UBERON:0001008), Reproductive system (UBERON:0000990), Respiratory system (UBERON:0001004) and Skeletal system (UBERON:0001434). We then retrieved all the sub-structures under these high-level terms down to the leaves of the ontology. We randomly picked five terms of the final lists of structures to verify manually that they indeed corresponded to the appropriate anatomical systems. We did not find any false positives during this process.

Similarly to the lists of nervous system structures, we retained in these lists only anatomical structures showing expression of at least 5 genes.

### Rate of sequence evolution

We retrieved the rate of non-synonymous substitutions *d*_N_ and the rate of synonymous substitutions *d*_S_ for mouse genes from Ensembl release 79 (Hubbard et al. 2009) using BioMart (Smedley et al. 2009). The *d*_N_ and *d*_S_ values were calculated pairwise using one-to-one orthologs in rat (see http://www.ensembl.org/info/genome/compara/homology_method.html#dnds).

### Gene Ontology

We retrieved genes annotated to the Gene Ontology category “metabolic process” (GO:0008152) and its sub-categories from the UniProtKB/Swiss-Prot database (The UniProt Consortium 2015), using the following URL: http://www.uniprot.org/uniprot/?query=reviewed:yes+organism:%22Mus%20musculus%20(Mouse)%20[10090]%22+go:8152 (queried on Aug 2^nd^, 2016). We performed a similar query to retrieve genes annotated to the category “membrane” (GO:0016020; Supplementary Text).

### Protein complexes

We obtained the precise annotation of number of subunits in protein complexes from manually curated information in the UniprotKB/Swiss-Prot database (The UniProt Consortium 2015). We downloaded data on July 21^st^, 2016 using the following URL: http://ebi4.uniprot.org/uniprot/?sort=&desc=&compress=no&query=&fil=reviewed:yesANDorganism:“Musmusculus(Mouse)[10090]”&force=no&preview=true&format=tab&columns=id,genes,comment(SUBUNIT). We used regular expressions in a Perl script (available on our GitHub repository) to extract the free-text annotation about involvement in protein complexes in the “SUBUNIT” annotation field. We divided genes into the following categories: monomers (524 genes), homo-multimers (1,936 genes), hetero-dimers (746 genes), hetero-multimers with more than two subunits (e.g., hetero-trimers; 327 genes), and all other complexes that are not described precisely enough to be classified automatically (1,075 genes). The lists of genes in the different categories are available as supplementary material on our GitHub repository (mouse_complexes.zip). If a gene was annotated in multiple categories, we kept it only in the “highest” category, following this order: hetero-multimer > hetero-dimer > uncharacterized complexes > homo-multimers > monomers.

### Connectivity

We retrieved the numbers of direct neighbors of genes in the mouse protein-protein interactions network from the OGEE database. We downloaded the file connectivity.txt.gz at this link: http://ogeedb.embl.de/#download, on July 7^th^, 2016.

### Akashi’s test

Selection for translational accuracy was tested using Akashi’s test (Akashi 1994; Drummond and Wilke 2008), following the procedure described at http://drummond.openwetware.org/Akashi’s_Test.html. Alignments of mouse and rat protein-coding genes were retrieved from Ensembl using the Perl API. Sites with the same amino acid at the aligned position in mouse and rat sequences were designated conserved. Optimal codons in mouse were taken from Drummond and Wilke (2008). Laplace smoothing was applied to contingency tables in order to remove problems with counts of zero. The outputs of the test are: (i) a *Z* score, which assesses how likely the association in a gene sequence between conserved sites and preferred codons is to have occurred by chance (significance), and (ii) a *Psi* score that assesses how strong is the association between preferred codons and conserved sites, which is computed as an odds ratio.

### Translation rates

We downloaded the mean of the typical decoding rates (MTDR) index for mouse genes in embryonic stem cells, embryonic fibroblasts and neutrophils from http://www.cs.tau.ac.il/~tamirtul/MTDR/MTDR_ORF_values/ (Dana and Tuller 2014). The MTDR index represents the geometrical mean of the typical nominal translation rates of codons of a gene, estimated from ribosome profiling data, after filtering biases and the effects of phenomena such as ribosomal traffic jams and translational pauses.

### False discovery rates

A false discovery rate of 10% was used to reported anatomical structures showing expression enrichment. For following analyses, where we disentangle the multiple factors associated with duplicate retention rates, we did not find a convenient way to correct for multiple testing. When enough independent tests of similar nature are performed, it is possible to estimate false discovery rates, but all our tests are not independent. Nonetheless, to give a rough estimate of the false discovery rate in these analyses, we collected all *p*-values generated for the linear regressions of the bin analyses in this paper (51 *p*-values). There was a clear excess of small *p*-values among them, indicating the presence of genuine signal (Figure S15). Using this list of *p*-values, we estimated that at a *p*-value threshold of 5%, the false discovery rate was well-controlled, at 10.2% using the FDR method (Benjamini and Hochberg 1995), or 3.4% using the q-value method (Storey and Tibshirani 2003).

## Acknowledgements

We thank Frederic Bastian for help with Bgee data retrieval; members of the Robinson-Rechavi lab, Allan Drummond and Aoife McLysaght for helpful discussions; Adrian Alexa for help in the adaptation of the topGO package, and the ZFIN and GXD teams for providing *in situ* hybridization data and help with data retrieval. Part of the computations were performed at the Vital-IT (http://www.vital-it.ch) Center for high-performance computing of the SIB Swiss Institute of Bioinformatics. We acknowledge funding from Etat de Vaud and Swiss National Science Foundation grant 31003A_153341 to MRR, Marie Curie IOF 273290 to JR, and Swiss Institute of Bioinformatics for support of the Bgee database.

## Author contributions

JR made the original observation; JR and JL designed the detailed study with input from MRR; JR and JL analyzed the data; JR wrote the paper with input from JL and MRR.

## Supplementary figures

Figure S1:

Illustration of the tree topologies targeted in this paper to identify duplicate and singleton genes. See “identification of duplicates and singletons” in the Materials and Methods for explanations complementing this figure. Black circles represent speciation nodes and red squares duplication nodes. Lineages in dotted lines represent lineages that were allowed in the selected topologies, but could be absent in some trees. Lineages names in red represent the target species (zebrafish or mouse). Internal node annotations represent the targeted taxonomical levels.

A: 3R ohnologs in zebrafish.

B: 3R singletons in zebrafish.

C: mouse orthologs of zebrafish 3R ohnologs.

D: mouse orthologs of zebrafish 3R singletons.

E: 2R ohnologs in mouse

F: 2R singletons in mouse

G: Small-scale duplicates in zebrafish. Paralogs with 100% or <10% sequence identity were filtered out.

H: Small-scale duplicates in mouse. Paralogs with 100% or <10% sequence identity were filtered out.

Figure S2:

(A) Proportion of mouse orthologs of zebrafish 3R ohnologs among genes expressed in the different tissues sampled in the GSE10246 microarray experiment. Legend as Figure 1.

(B) Boxplot of the proportion of human orthologs of zebrafish 3R ohnologs among genes expressed in the different tissues sampled in the GTEx RNA-seq experiment.

(C) Proportion of mouse orthologs of zebrafish 3R ohnologs among genes expressed in the different tissues sampled in the GSE16496 microarray experiment.

Figure S3:

Similar to Figure 2B, but nervous system genes are those that were not expressed in any other non-nervous structure.

Figure S4:

A: Spearman’s correlation coefficient between *d*_N_ and the expression levels in the different tissues sampled in the GSE3594 microarray experiment. Tissues are ranked based on the average correlation coefficient across biological replicates, represented with different dots. Green color represents nervous-system tissues and purple represents non-nervous-system tissues. “Mean expression” and “maximum expression” represent the correlation between *d*_N_ and the mean or maximum expression levels across all samples. “Mean nervous system expression” and “maximum nervous system expression” represent the correlation between *d*_N_ and the mean or maximum expression levels across all samples from nervous system tissues. B: Similar to (A), but using samples from the GSE16496 microarray experiment including only nervous tissues. “Mean nervous system expression*” and “maximum nervous system expression*” represent the correlation between *d*_N_ and the mean or maximum expression levels across all samples from nervous system tissues, excluding samples from the three nervous tissues displaying lower correlation than other: retina, pineal body and pituitary gland.

Figure S5:

Scatterplots illustrating, in mouse, the pairwise relationships between the gene properties used in this paper. Spearman’s correlation coefficients (rho) and *p*-values are indicated in the top-right corner of the figures. Loess regression lines are plotted in red.

A: relation between *d*_N_ and mean expression level across nervous system samples from the GSE16496 microarray experiment.

B: relation between *d*_N_ and maximum expression level across nervous system samples from the GSE16496 microarray experiment.

C: relation between *d*_N_ and *d*_S_.

D: relation between *d*_S_ and maximum nervous system expression level.

E: relation between *d*_N_ and Akashi’s test *Psi* score.

F: relation between *Psi* and maximum nervous system expression level.

G: relation between *d*_N_ and connectivity (number of protein-protein interactions).

H: relation between *Psi* and connectivity.

I: relation between connectivity and maximum nervous system expression level.

Figure S6:

A: Relation between proportion of mouse orthologs of zebrafish 3R ohnologs and expression levels summaries, mean and maximum nervous system expression. Only nervous system genes were used for this analysis. Genes were split into 10 equal-sized bins of expression levels, and the median expression level of each bin was plotted on the x-axis (in log_2_ scale). The analysis using mean nervous system expression is plotted in dashed line and open circles, while the analysis using maximum nervous system expression is plotted in solid line and plain circles.

B: Similar to A, using maximum nervous system expression. The analysis including the 20% genes with highest *d*_N_ is plotted in blue and squares, while the analysis including 20% genes with lowest *d*_N_ is plotted in red and triangles.

C: Similar to Figure 3C. The analysis including the 20% genes with highest maximum nervous system expression is plotted in red and squares, while the analysis including the 20% genes with lowest maximum nervous system expression is plotted in blue and triangles.

Figure S7:

A: Spearman’s correlation coefficient between *d*_S_ and the expression levels in the different tissues sampled in the GSE3594 microarray experiment. Legend similar to Figure S4.

B: Relation between proportion of mouse orthologs of zebrafish 3R ohnologs and rate of synonymous substitution. Legend similar to Figure 3A and B.

Figure S8:

Relation between Akashi’s test *Psi* score and translation rates across three different cell types in mouse. Legend similar to Figure S5.

A: neutrophils

B: embryonic fibroblasts

C: embryonic stem cells

Figure S9:

A: Proportion of mouse orthologs of zebrafish 3R ohnologs for members of different complexes subtypes, classified using UniProtKB/Swiss-Prot data. The number of genes in each category is indicated at the bottom of each box. Green bars represent nervous system genes and purple bars represent non-nervous system genes.

B: Relation between proportion of mouse orthologs of zebrafish 3R ohnologs and *d*_N_, controlling for complex membership. Legend similar to Figure 3C. Genes were divided into different groups based on their membership in protein complexes, using data from the UniProtKB/Swiss-Prot database.

Figure S10:

Proportion of mouse orthologs of zebrafish 3R ohnologs for metabolic process and non-metabolic process genes, based on Gene Ontology annotation. Legend similar to Figure S9A.

Figure S11:

A: Comparison of connectivity for mouse orthologs of zebrafish 3R ohnologs or singletons, depending on their nervous system expression. Legend similar to Figure 2.

B: Relation between proportion of mouse orthologs of zebrafish 3R ohnologs and *d*_N_, controlling for connectivity. Legend similar to Figure 3.

C: Relation between proportion of mouse orthologs of zebrafish 3R ohnologs and connectivity, controlling for *d*_N_. Legend similar to Figure 3.

D: Relation between proportion of mouse orthologs of zebrafish 3R ohnologs and connectivity, controlling for complex membership. Legend similar to Figure S9B.

Figure S12:

Relation between the proportion of small-scale duplicates in mouse, and nervous system expression, *d*_N_, and *Psi* score.

A: Similar as Figure 2.

B and C: Similar to Figure 3C and D, but including all genes.

Figure S13:

A: Relation between proportion of mouse small-scale duplicates and expression levels summaries, mean and maximum expression across all samples from the GSE3594 microarray experiment. Legend similar to Figure S6A. The analysis using mean expression is plotted in plain line and plain circles, while the analysis using maximum expression is plotted in dashed line and open circles.

B and C: Similar to Figure S6B and C for mouse small-scale duplicates, using mean expression level.

Figure S14:

Proportion of mouse small-scale duplicates for members of different complexes subtypes. Legend similar to Figure S9.

Figure S15:

Histogram of *p*-values from 51 linear regressions performed in Figures 3, S6, S7, S9, S11, S12, S13 and S17. A dashed red line indicates the 5% significance threshold used in this study.

Figure S16 (Supplementary Text):

Histogram of *p*-values from 10,000 Wilcoxon tests obtained by comparing the *d*_N_ values of nervous system orthologs of ohnologs and singletons, after randomly resampling the set of nervous system genes to the same size as the set of non-nervous system genes. A dashed red line indicates p=0.05, and a plain red line indicates p=0.41 (the p-value obtained from the Wilcoxon test between non-nervous system orthologs of ohnologs and singletons, see Figure 2B).

Figure S17 (Supplementary Text):

A: Proportion of mouse orthologs of zebrafish 3R ohnologs for members of different complexes subtypes, classified using data from the CORUM database. Similar to Figure S9.

B: Proportion of mouse small-scale duplicates for members of different complexes subtypes. Similar to Figure S14.

Figure S18 (Supplementary Text):

A: Proportion of mouse orthologs of zebrafish 3R ohnologs for genes encoding membrane proteins or not, based on Gene Ontology annotation. Legend similar to Figure S9A.

B: Relation between proportion of mouse orthologs of zebrafish 3R ohnologs and *d*_N_, controlling for the effect of encoding a membrane protein. Legend similar to Figure 3.

## Supplementary tables

Table S1:

Zebrafish anatomical structures showing a significant enrichment in expression of zebrafish 3R ohnologs. Similar to Table 1, but all structures are shown, even non-significant ones.

Table S2:

Zebrafish anatomical structures showing a significant depletion in expression of 3R singletons.

Table S3:

Same as Table S1 but using an independent list of zebrafish 3R ohnologs obtained from (Braasch et al. 2016).

Table S4:

Same as Table S2 but using an independent list of zebrafish 3R singletons obtained from (Braasch et al. 2016).

Table S5:

Mouse anatomical structures showing a significant enrichment in expression of orthologs of zebrafish 3R ohnologs.

Table S6:

Mouse anatomical structures showing a depletion in expression of orthologs of zebrafish 3R singletons.

Table S7:

Mouse anatomical structures showing a significant enrichment in expression of 2R ohnologs.

Table S8:

Same as Table S7 but using an independent list of mouse 2R ohnologs obtained from (Singh et al. 2015). The three lists of 2R ohnologs, calculated with different levels of stringency are available at http://ohnologs.curie.fr/cgi-bin/BrowsePage.cgi?org=mouse. Here, the most stringent list was used, but the results were similar with the intermediate and relaxed lists.

Table S9:

Mouse anatomical structures showing a significant depletion in expression of 2R singletons.

Table S10:

Mouse anatomical structures showing a significant depletion in expression of small-scale duplicates.

Table S11:

Same as Table S10 but using an independent list of rodent-specific small-duplicate genes obtained from (Farre and Alba 2010).

Table S12:

Mouse anatomical structures showing a significant enrichment in expression of slowly evolving genes (the 10% lowest *d*_N_ values based on mouse-rat comparisons; FDR < 10%).

Table S13:

Mouse anatomical structures showing a significant enrichment in expression of genes with the 10% highest *Psi* score from Akashi’s test.

